# Quaternary structure evaluation tool for protein assemblies

**DOI:** 10.1101/224196

**Authors:** Selcuk Korkmaz, Jose M. Duarte, Andreas Prlić, Dincer Goksuluk, Gokmen Zararsiz, Osman Saracbasi, Stephen K. Burley, Peter W. Rose

## Abstract

The Protein Data Bank (PDB) is the single worldwide archive of experimentally-determined three-dimensional (3D) structures of proteins and nucleic acids. As of January 2017, the PDB housed more than 125,000 structures and was growing by more than 11,000 structures annually. Since the 3D structure of a protein is vital to understand the mechanisms of biological processes, diseases, and drug design, correct oligomeric assembly information is of critical importance. For example, it makes a difference if the protein is normally a dimer and not a monomer or a trimer or a tetramer or a hexamer in nature. Unfortunately, the biologically relevant oligomeric form of a 3D structure is not directly obtainable by X-ray crystallography. Instead, this information may be provided by the PDB Depositor as metadata coming from additional experiments, be inferred by sequence-sequence comparisons with similar proteins of known oligomeric state, or predicted using software, such as PISA (Proteins, Interfaces, Structures and Assemblies) or EPPIC (Evolutionary Protein Protein Interface Classifier). Despite significant efforts by professional PDB Biocurators during data deposition, there remain a number of structures in the archive with incorrect quaternary structure descriptions (or annotations). Further investigation is, therefore, needed to evaluate the correctness of quaternary structure annotations. In this study, we aim to identify the most probable oligomeric states for proteins represented in the PDB. Our approach evaluated the performance of four independent prediction methods, including text mining of primary publications, inference from homologous protein structures, and two computational methods (PISA and EPPIC). Aggregating predictions to give consensus results outperformed all four of the independent prediction methods, yielding 86% correct, 9% incorrect, and 5% inconclusive predictions, when tested with a well-curated benchmark dataset. We have developed a freely-available web-based tool to make this approach accessible to researchers and PDB Biocurators (http://quatstruct.rcsb.org).

## Introduction

The Protein Data Bank (PDB, pdb.org) [1] provides detailed information about the three-dimensional (3D) structures of biological macromolecules, including proteins and nucleic acids. The PDB was established in 1971 with only 7 X-ray crystal structures of proteins and now contains more than 125,000 structures (as of January 2017). Today, the PDB archive is managed by the international Worldwide Protein Data Bank (wwPDB, wwpdb.org) partnership [2], which includes the RCSB Protein Data Bank (RCSB PDB, rcsb.org) [1], the Protein Data Bank in Europe (PDBe, pdbe.org), Protein Data Bank Japan (PDBj, pdbj.org), and BioMagResBank (BMRB, bmrb.org). The majority (~90%) PDB structures were determined by X-ray crystallography. This experimental method yields 3D atomic level structures of the so-called asymmetric unit (Fig 1A), which is the repeating unit that makes up the crystal (Fig 1B). Knowledge of the 3D structure of the asymmetric unit and intermolecular interactions among asymmetric units does not provide sufficient information to reveal conclusively the oligomeric structures of protein assemblies, because is often not possible to distinguish biologically relevant intermolecular contacts from contacts that merely stabilize the crystal lattice.

**Fig 1.**
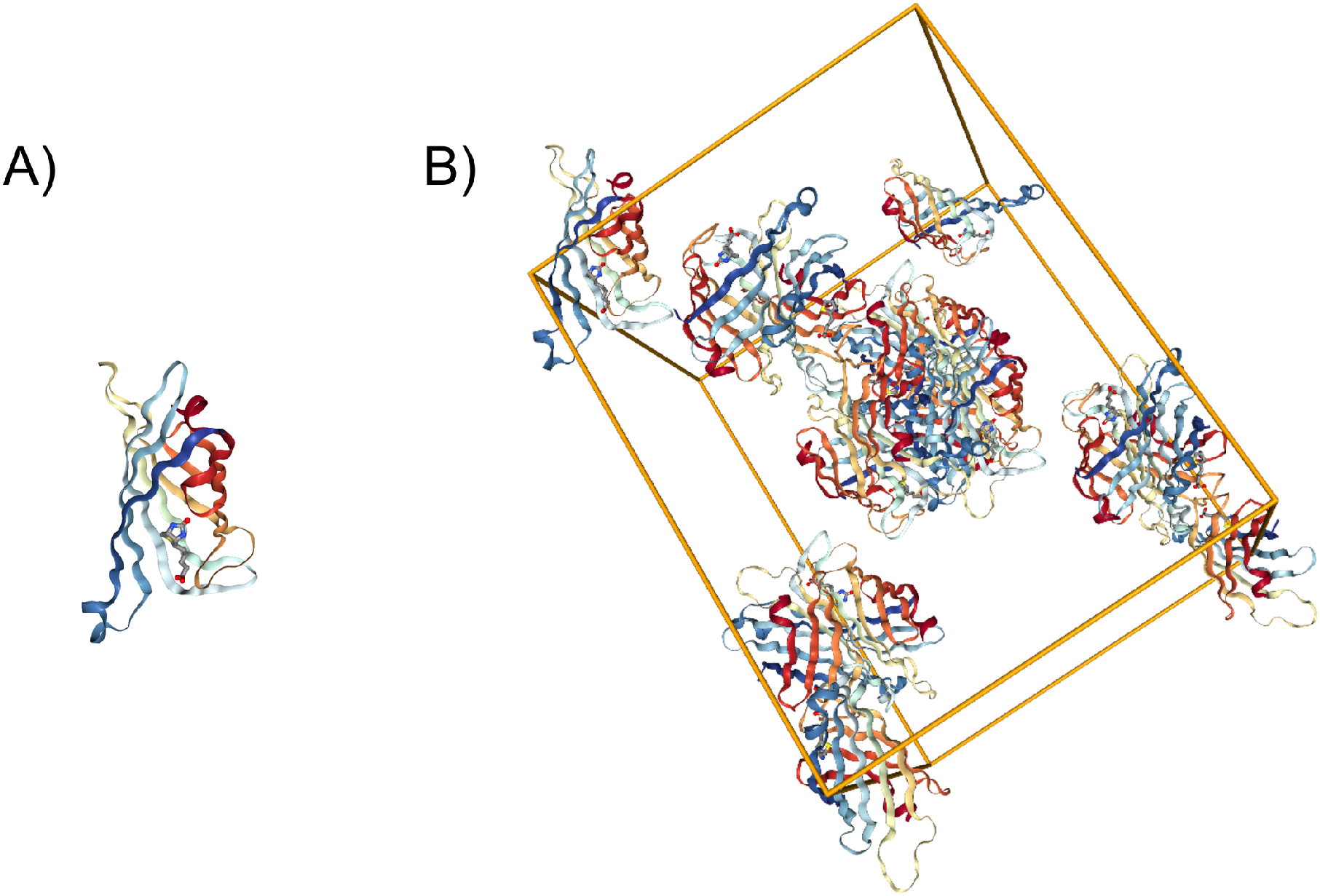
3D structure information for PDB entry 1STP. A) Monomeric asymmetric unit. B) Unit cell containing asymmetric units within the crystal lattice.

Many proteins form structurally well-characterized thermodynamically stable multimeric complexes, which are important for biological function [e.g., hemoglobin occurs in nature a heterotetramer (A2B2) with a cyclic (C2) symmetry and dihedral (D2) pseudo-symmetry] [3]. Experimental methods, such as size exclusion chromatography or analytical ultracentrifugation are sometimes required to ascertain the correct oligomerization state for a protein structure determined by X-ray crystallography. Alternatively, correct oligomeric state information may be inferred by comparison with better characterized homologous proteins or be provided by the PDB Depositor as metadata. It can also be predicted using computational methods, such as PISA (Proteins, Interfaces, Structures, and Assemblies) [4] or EPPIC (Evolutionary Protein-Protein Interface Classifier) [5]. Since the PDB was established in 1971, oligomeric state information has been obtained from Depositors or predicted by PQS [6] and more recently with PISA. Although experimental evidence for oligomeric state was not a mandatory data item in legacy PDB deposition systems, collection of experimental evidence has been improved in the new wwPDB OneDep global deposition, biocuration, and validation system [7].

Quaternary structures of proteins can be characterized by two main descriptors that define their oligomeric states: stoichiometry and symmetry. Stoichiometry describes the composition of the assembly in terms of subunit number and composition. There are several widely used methods for determining the stoichiometry of protein complexes, including size exclusion chromatography [8], analytical ultracentrifugation [9], and gel-electrophoresis [10, 11]. Protein assembly stoichiometry is described using a composition formula. Typically, an uppercase letter, such as A, B, C, etc., represents each type of different protein subunit in alphabetical order. (N.B.: These letters are not the same as the chain identifiers found in PDB archival entries.] The number of equivalent subunits is added as a coefficient next to each letter. For example the stoichiometry of the two-component human hemoglobin heterotetramer is represented as A2B2 (a dimer of heterodimers composed to two distinct polypeptide chains). Symmetry is another important feature of protein tertiary and quaternary structure [12] and plays a key role in understanding protein evolution and structure/function relationships [3], [12], [13], [14], [15], [16]. At the quaternary structure level, we characterize symmetry by the point group, a set of symmetry elements, whose symmetry axes go through a single point [12]. Most oligomeric protein structures (either homomeric or heteromeric) are symmetric macromolecules, which is probably a simple consequence of how subunits associate in solution without aggregating indefinitely [17] and can be classified using closed symmetry groups [18]. They are typically described as cyclic (i.e., C2, C3, C4,…), dihedral (i.e., D2, D3, D4,…), or cubic (tetrahedral, octahedral, icosahedral). Dihedral and cyclic symmetries are geometrically related: a structure with Dn symmetry can be constructed from n dimers with C2 or from two n-mers with Cn symmetry [19]. Additionally, helical symmetry is also a common open-symmetry encountered in protein structures.

The PDB archive grows by more than 11,000 structures annually. However, because of incomplete data and errors made during data entry, the oligomeric state annotations provided by the PDB are not always correct and reliable [20]. In spite of great efforts to improve the quality of the PDB archive, it has been reported that there are a significant number of PDB entries with incorrect quaternary structure annotations (Fig 2). Levy put the the error rate at ~14% [21], while more recently Baskaran et al. [22] reported a lower bound for the error rate of ~7%. Development of methods for accurate detection of incorrect annotations and assignment of most probable oligomeric state is, therefore, a matter of some urgency.

**Fig 2.**
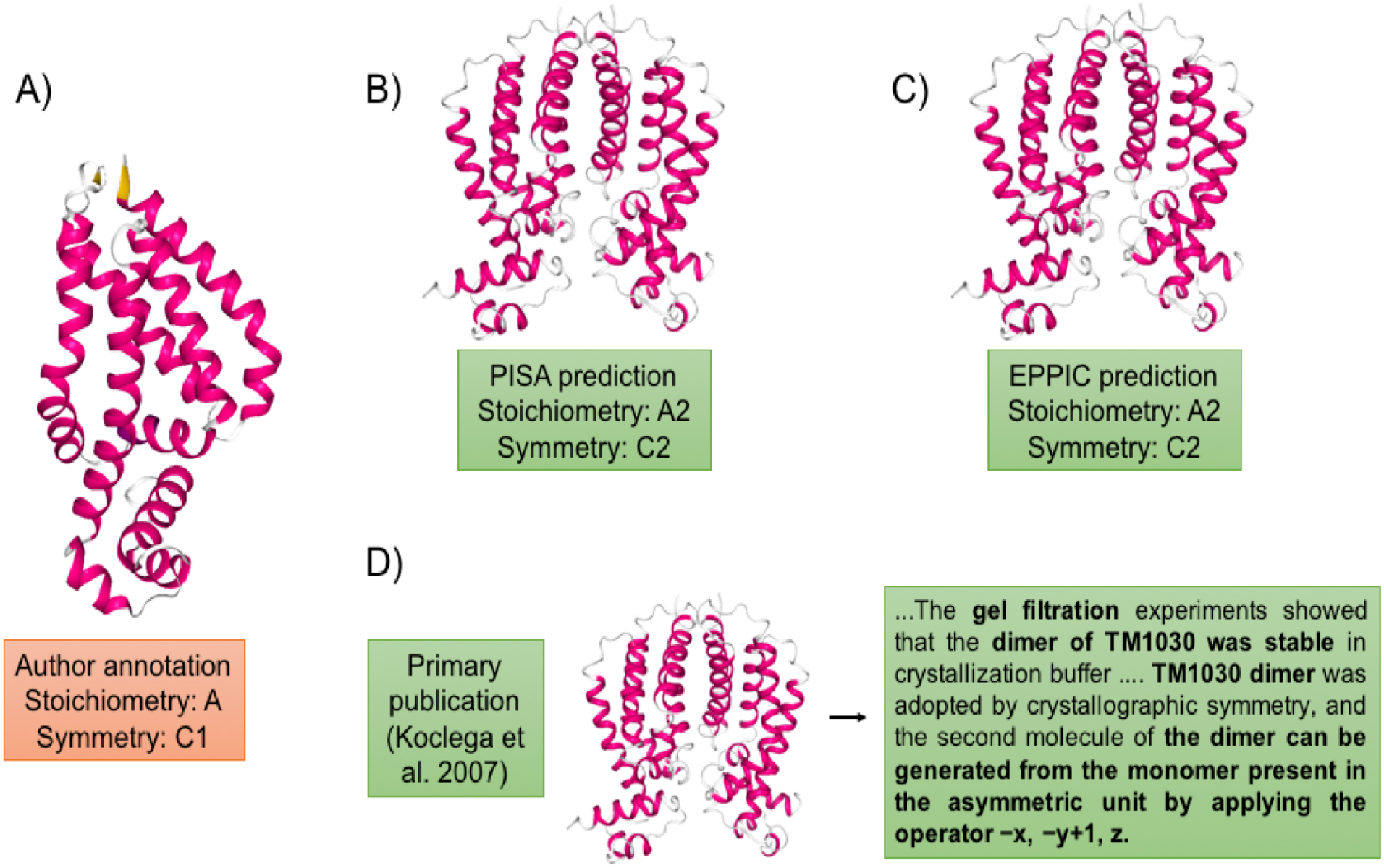
Quaternary structure annotations for PDB entry 1Z77. A) Incorrectly annotated as monomer during PDB deposition. Correctly annotated as a dimer by B) PISA, C) EPPIC, and D) the primary publication.

This study has two main objectives: (i) to enable identification of incorrect quaternary structure annotations in the PDB archive, and (ii) to enable assignment the most probable quaternary structure for such cases. To accomplish these goals, we evaluated four different methods for assessing quaternary structure annotations in the PDB. First, we took an evolutionarily approach by clustering proteins related by amino acid sequence and attributed to each member of a given cluster the oligomerization state found to be most prevalent with the cluster. Second, we took a text mining approach by searching through the primary citations of indivdiual PDB entries and extracting information about oligomeric state and experimental evidence thereof. Third and fourth, we took two independent computational approaches using PISA and EPPIC, respectively, to predict oligomeric states. We aggregated results from these methods to generate a consensus prediction for the most probable oligomeric state for each protein structure in the PDB. We tested this combined approach using a well-curated benchmark dataset. During the course of this effort, we developed an efficient approach to evaluate oligomeric states of protein structures in the PDB, which we have made freely available to both PDB Biocurators and researcher as a web-based tool.

## Materials and Methods

### Benchmark Dataset

We aggregated three previously published, manually curated, benchmark datasets to create a considerably larger combined benchmark dataset for this work. The Ponstingl dataset [23] contains 218 protein complexes, including 55 monomers, 88 dimers, 24 trimers, 38 tetramers, and 13 hexamers. The Bahadur dataset [24, 25] contains 266 PDB entries, including 144 monomers and 122 dimers. The Duarte dataset contains [5] 152 protein structures, including 78 monomers, 62 dimers, 2 trimers, 8 tetramers, 1 hexamer, and 1 dodecamer. After removing duplicate entries (Fig 3), our final combined benchmark dataset contains 543 biological macromolecules, including 248 monomers, 209 dimers, 26 trimers, 44 tetramers, 14 hexamers, 1 octamer and 1 dodecamer.

**Fig 3.**
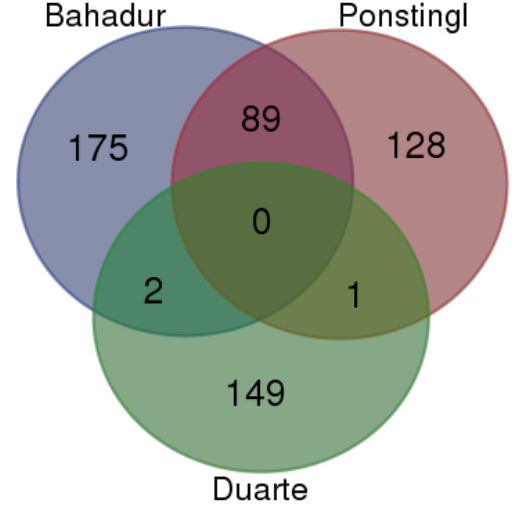
Overlapping structures between datasets. Most of the duplicates occcurred between the Ponstingl and the Bahadur datasets. These two datasets share 89 protein structures, whereas only 2 structures were found to be common between the Bahadur and the Duarte datasets, and only 1 structure was detected in both the Ponstingl and the Duarte datasets.

### Sequence Clustering

Since many protein chains in the PDB are similar at the level of sequence, we use this information to cluster polypeptide chains on the basis of amino acid sequence identity and assign a representative oligomeric state for each cluster based on a consistency score. For this purpose, we first constructed sequence clusters at various identity thresholds, including 95%, 90%, 70% and 40%. Clusters were calculated using the BLASTClust algorithm [26], which detects pairwise matches with the blastp algorithm [27] and then places each sequence in a certain cluster if the sequence matches that of at least one cluster member (Fig. 4).

**Fig 4.**
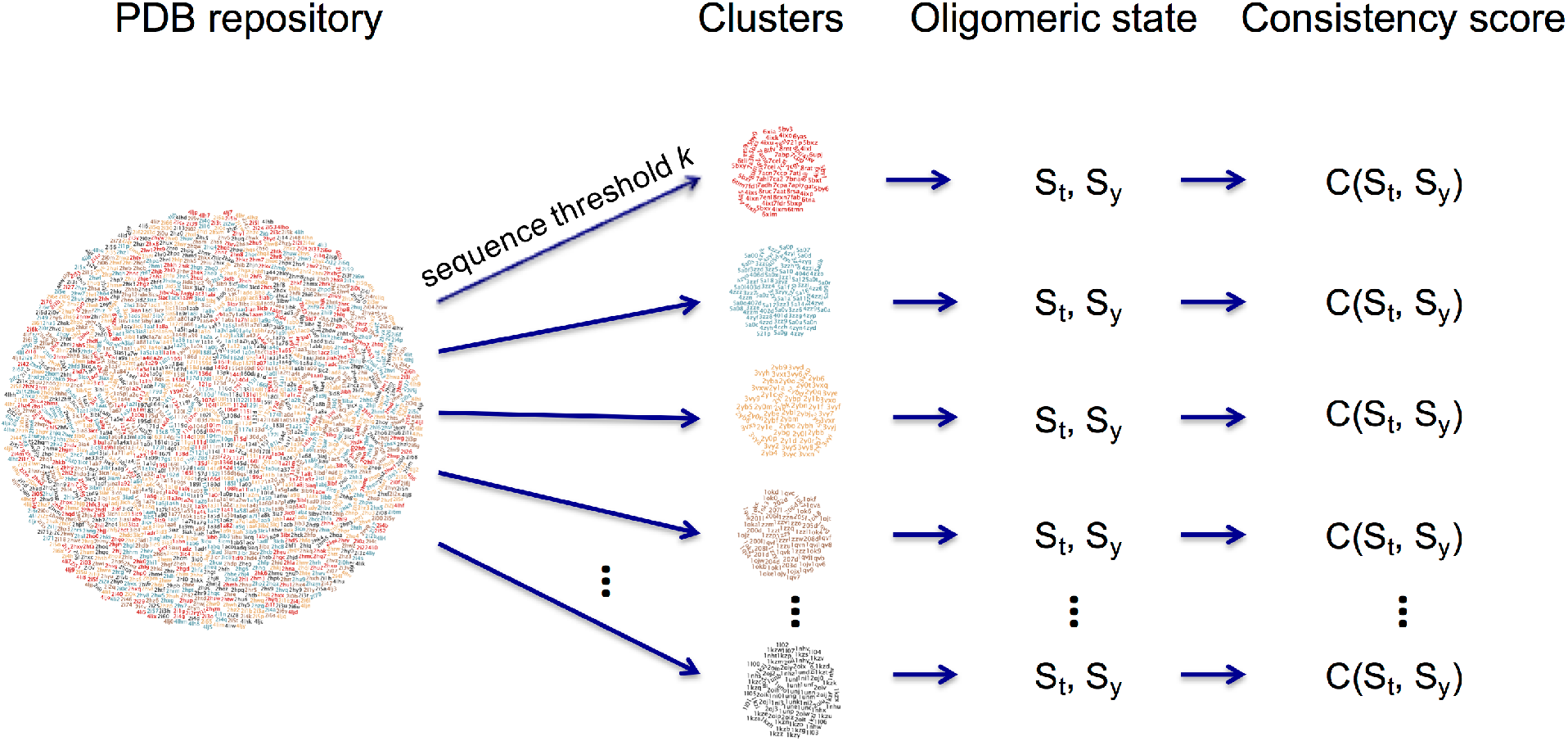
Sequence cluster construction and consistency score calculation. A consistency score is calculated based on the distribution of stoichiometry and symmetry values within each cluster. (S_t_: Stoichiometry, S_y_: Symmetry, C(S_t_, S_y_): consistency score for given S_t_ and S_y_).

A cluster is defined as a set of protein chains that are at least k% sequence identical to each other over 90% of the same length, and is associated with two discrete random variables:

- S_t_ represents stoichiometry and consists of a list (a,b,c,…,m) giving the number of copies of each unique molecule (e.g., A: monomer, A2: homodimer, A2B2: heterotetramer).
- S_y_ represents symmetry and takes on values such as C2 (cyclic), D4 (dihedral), H (helical).

For a certain sequence identity threshold k, a consistency score for a given stoichiometry t and symmetry y can be estimated by the joint probability of these two events:

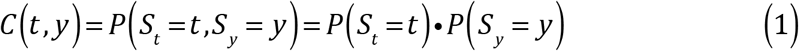

After consistency score calculation, we applied a binary decision rule based on the majority probability to predict a representative quaternary structure for a certain cluster. The maximum consistency score in a cluster must be greater than 0.5 to satisfy the majority rule and to predict a representative quaternary structure, otherwise the result is deemed inconclusive.

For the consistency score to be statistically meaningful, there must be minimum number of members in a cluster. We used our benchmark dataset to determine the minimum number of cluster members for different sequence cluster identity thresholds, including 40%, 70%, 90% and 95%. For this purpose, we selected different number of cluster sizes from (n=1,…,50) and predicted the most representative oligomeric state for each cluster using the consistency score described above. Then, we calculated the percentage of correct, incorrect, and inconclusive predictions for each minimum number of cluster members and for each sequence identity threshold. According to these results, each increment in the minimum number of cluster members reduces the number of incorrect results, but increases the number of inconclusive results. Hence, there is a trade-off between errors and inconclusive results. In Figs 5A-D, we plotted each prediction against cluster size and required at least 70% correct prediction for each sequence identity threshold. Lower sequence identities require more cluster members, while higher sequence identities require fewer. Based on our work with the benchmark dataset results, minimum cluster size should be 5 for 40% sequence identity threshold, yielding a 6% error rate and and 24% inconclusive rate. For 70%, 90%, and 95% sequence identities we found that the cluster size should be at least 3, yielding a 4% error rate and a 26% inconclusive rate.

**Fig 5.**
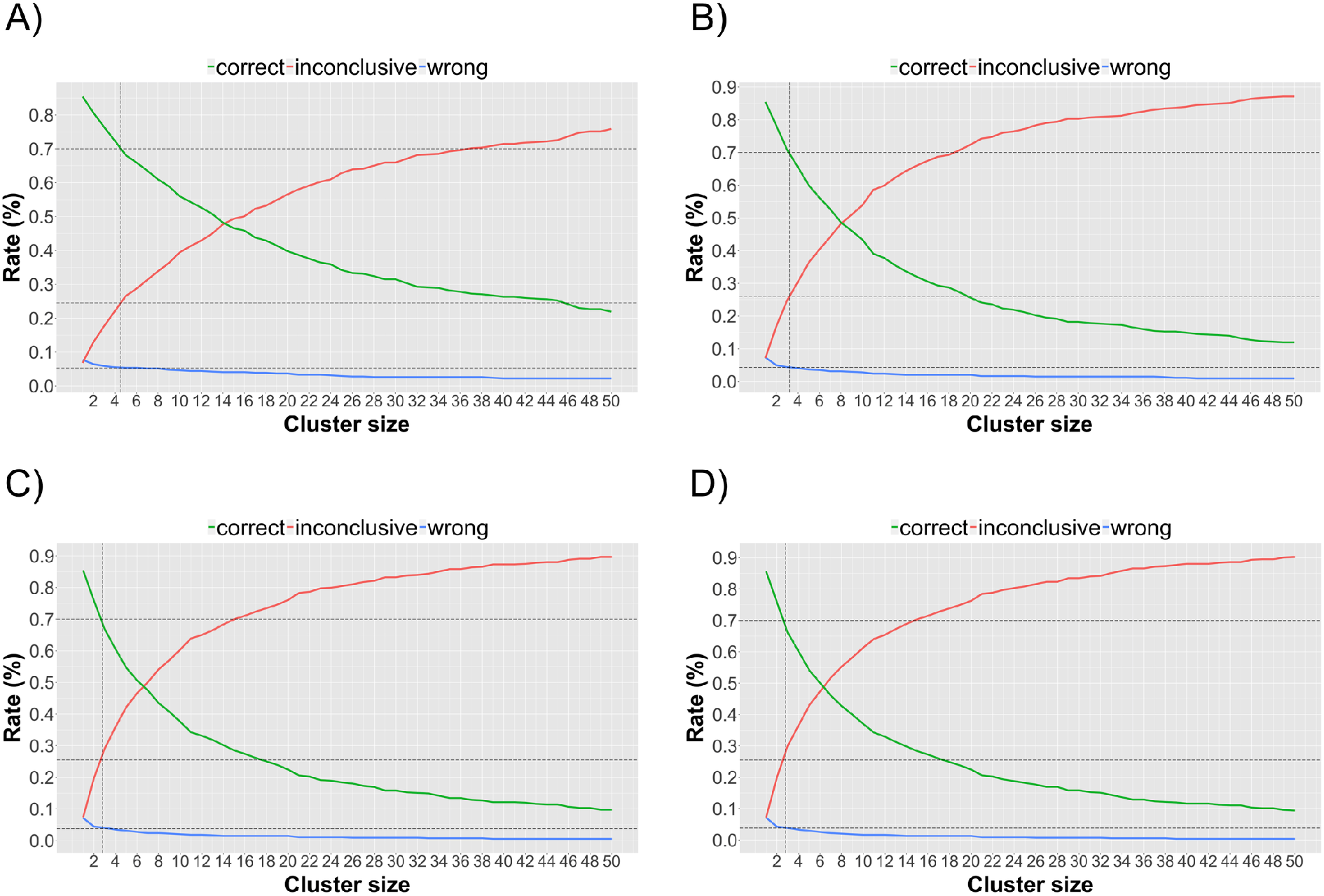
Cluster size vs. correct, incorrect, and inconclusive rates. A) For 40% sequence identity. B) For 70% sequence identity. C) For 90% sequence identity. D) For 95% sequence identity.

### Text Mining

Nearly 82% of the protein structures archived in the PDB have an associated primary article as of January 2017. Using CrossRef TDM (text and data mining) services (http://tdmsupport.crossref.org/), we were able to extract information regarding oligomeric state and supporting experimental evidence from 8,600 primary publications, describing nearly 32,000 PDB entries.

We first split the full-text article into sentences. Then, we identified sentences containing oligomeric state information using a keyword list (monomer, dimer, trimer, etc., see Table S1 for the full keyword list). Some of these sentences proved misleading or irrelevant. For example, some sentences described the asymmetric unit not quaternary structure, and some sentences refer to protein structures other than the one of interest. We, therefore, used a machine-learning approach to eliminate non-relevant sentences by classifying sentences as quaternary structure relevant (positive) or irrelevant (negative). To do so with traditional machine learning algorithms that require numerical inputs, each sentence had to be tokenized into words. Then, we converted each word to numerical values using the term frequency-inverse document frequency (tf-idf) method [28] to create a numerical data matrix. This method reflects how important a word is to a document in a corpus using the following formula:

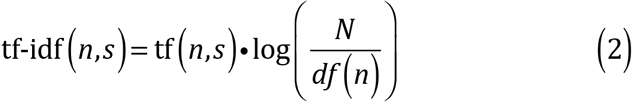

were tf(n,s) represents the frequency of word n in sentence s, df(n) represents the number of sentences containing word n, and N is the total number of sentences. To improve the effectiveness of the tf-idf method, all words in each sentence were converted to lower case, with extra spaces and internal punctuation marks removed. Finally, we calculated the tf-idf score for each word in each sentence and created a data matrix for the training procedure (wherein each row represented a single sentence and each column represented a unique word). To avoid the high-dimensional data matrix, we mapped the each column (i.e. features) to a hash-table by using a hash function. In this study, we used the murmurhash3 hash function, proposed by Weinberger et al. [29]. After applying the hashing function, we used two machine-learning algorithms, support vector machines (SVM) [30] and boosted logistic regression (BLR) [31–34], to classify each sentence in a paper as positive and negative.

A similar approach was used to search sentences for experimental evidence of oligomeric state. An experimental evidence keyword list was used for this task (see S2 Table for the full list). After oligomeric state keyword filtering, a second filtering was applied based on these keywords. Finally, after two filtering procedures, the quaternary structure prediction for each PDB entry was made based on the majority probability of remaining oligomeric state keywords. The text mining result was deeemed inconclusive for a particular PDB entry, if machine learning algorithms failed to detect any quaternary structure related sentences and supporting experimental evidence. A general workflow of our text mining approach can be found in Fig 6.

**Fig 6.**
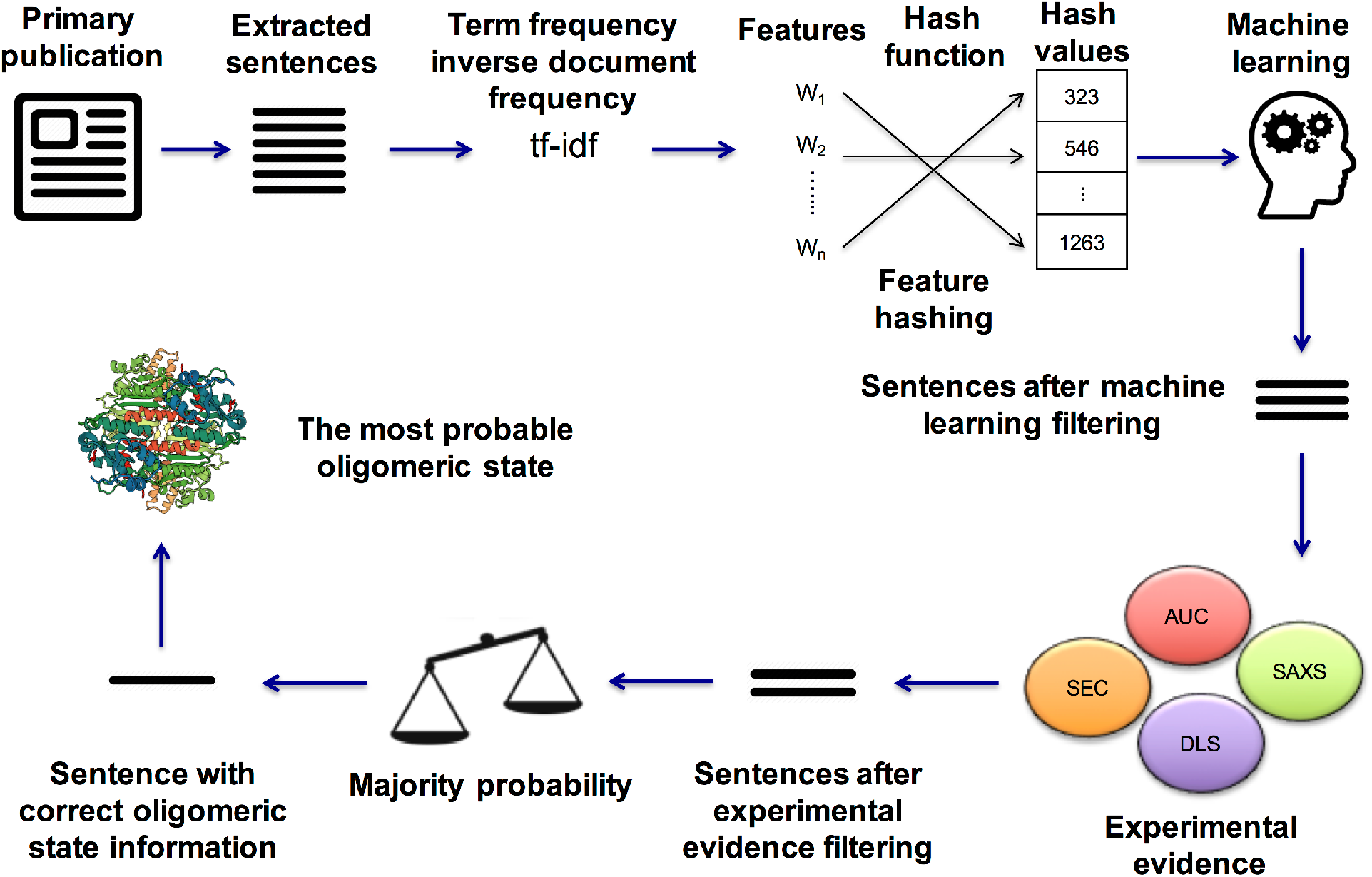
General workflow of the text mining approach. After a publication upload or download, the term frequency-inverse document frequency (tf-idf) method is used to create numerical data matrix from words, a hash table is created using the hash function, extracted sentences are classified using machine learning alorithms, remaining sentences are searched for experimental evidence and oligomeric state information is determined by using a majority rule.

To train and test our machine learning algorithms for the text mining approach, we created a dataset using PDB primary papers. For this task, first, we extracted sentences from the papers using the keyword list in the S1 Table. Then, the keyword list in the S2 Table was used to split these sentences into two classes. The sentences, which match to this second keyword list, were used to create the positive dataset of 5500 positive sentences and the remaining sentences were used to create the negative dataset of 5500 negative sentences. Next, the dataset was split as 80% training and 20% test set. A grid search was used with 10-fold cross-validation to select optimal parameters in the training set. Two parameters optimized for the SVM algorithm were sigma = 0.013 and cost = 4, and the optimal number of boosting iterations found as 101 for BLR algorithm. Identical parameters were used for the test set to confirm that both datasets were on the same scale and homoscedastic relative to each other. Finally, we tested model performance on the test set (Table 1). SVM performed better than BLR in terms of accuracy, kappa, area under the ROC curve (AUC), sensitivity, negative predictive value, F1 score, and Matthews correlation coefficient. Conversely, BLR showed better specificity and positive predictive value results, suggesting that it predicts positive sentences slightly better than SVM.

**Table 1.**
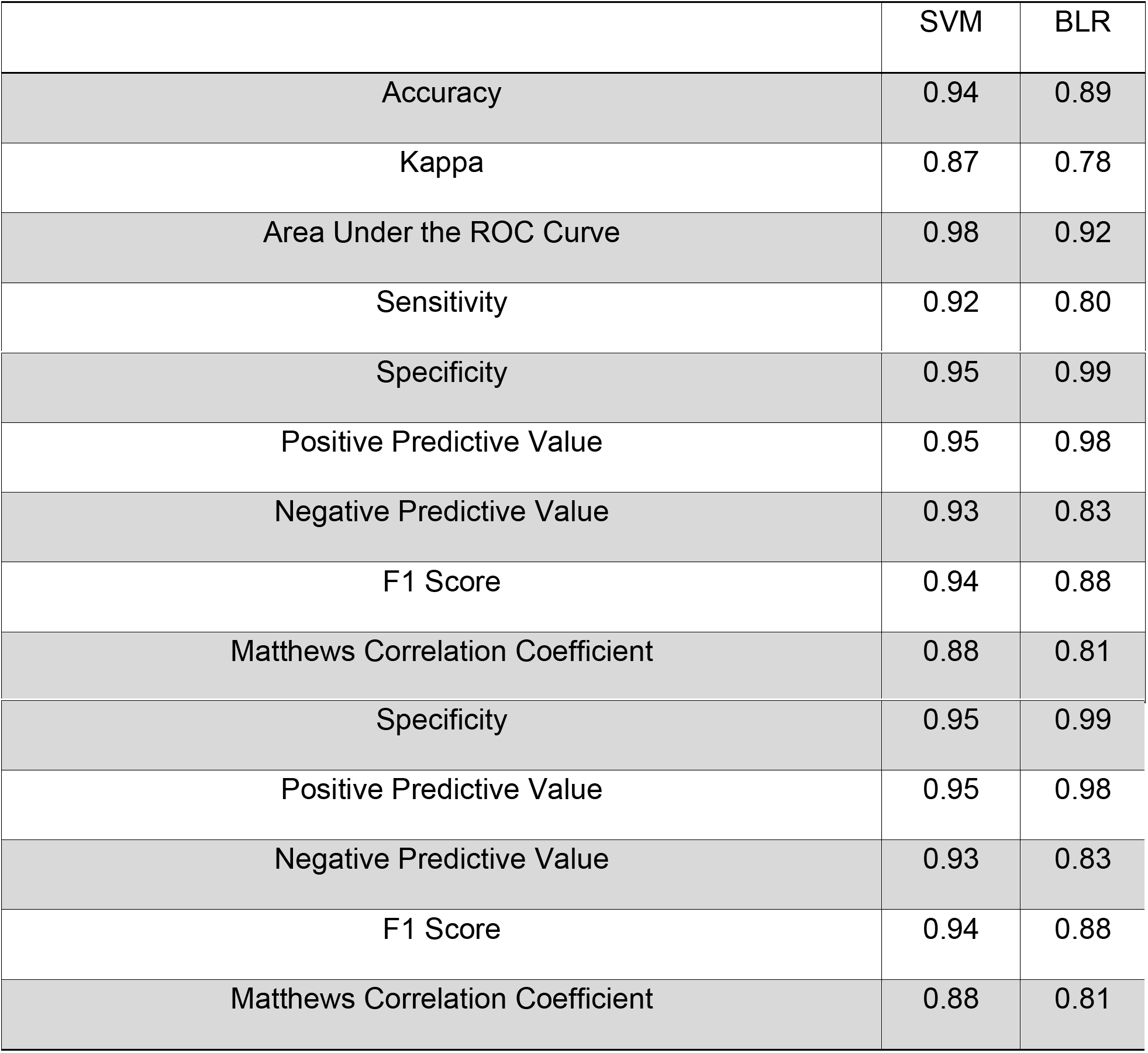
Test set (20% of the original data) performances of the machine learning algorithms. SVM outperforms BLR on a number of performance measures, whereas BLR has slightly better specificity and positive predictive value results.

|  | SVM | BLR |
| --- | --- | --- |
| Accuracy | 0.94 | 0.89 |
| Kappa | 0.87 | 0.78 |
| Area Under the ROC Curve | 0.98 | 0.92 |
| Sensitivity | 0.92 | 0.80 |
| Specificity | 0.95 | 0.99 |
| Positive Predictive Value | 0.95 | 0.98 |
| Negative Predictive Value | 0.93 | 0.83 |
| F1 Score | 0.94 | 0.88 |
| Matthews Correlation Coefficient | 0.88 | 0.81 |

### PISA Prediction

Following successful crystallographic structure determination efforts are made to identify biologically relevant intermolecular interactions within the crystal [35], and distinguish them from intermolcular contacts that simply stabilize the crystal lattice. The PISA program, developed by Krissinel and Henrick [4], uses a quantitative approach to address this problem [35]. The stability of an oligomeric structure is a function of free energy formation, solvation energy gain, interface area, hydrogen bonds, salt-bridges across the interface, and hydrophobic specificity [4]. PISA uses these properties to analyze protein structures and predict possible stable oligomeric states. Following successful evaluation (i.e., 90% accuracy [20]) using the Ponstingl et al. [23] benchmark data in 2007, PISA was deployed as a web server at the European Bioinformatics Institute (EBI) [35]. Soon thereafter, it was adopted as a quaternary structure validation and annotation tool for PDB archival depositions.

PISA can be accessed through a web service (http://www.ebi.ac.uk/) or as a standalone program (http://www.ccp4.ac.uk/pisa/) from Collaborative Computational Project No. 4 or CCP4 (http://www.ccp4.ac.uk/). The software provides broad information about assemblies, interfaces, and monomers, and gives information regarding possible oligomeric states, such as stoichiometry, solvent accessible surface area (ASA), buried surface area (BSA), and Gibbs free energy of dissociation score, which represents the free energy difference between the associated and dissociated states [36]. Possible oligomeric states yielding positive and negative Gibbs free energy of dissociation score are considered to be chemically stable and unstable quaternary structures, respectively [37]. Those with borderline values are deemed indeterminate.

In this study, we used the command line version of PISA to generate XML files for each possible oligomeric state, which include rotation/translation operators for each of the chains together with ASA, BSA, Gibbs free energy of dissociation score, entropy, internal energy, and macromolecular size. The PISA XML files were then parsed to extract rotation/translation operators and chains with which to build up the atomic coordinates of each possible oligomeric state. Finally, we used BioJava [38] to characterize each of the possible oligomeric states in terms of stoichiometry and symmetry (Fig 7).

**Fig 7.**
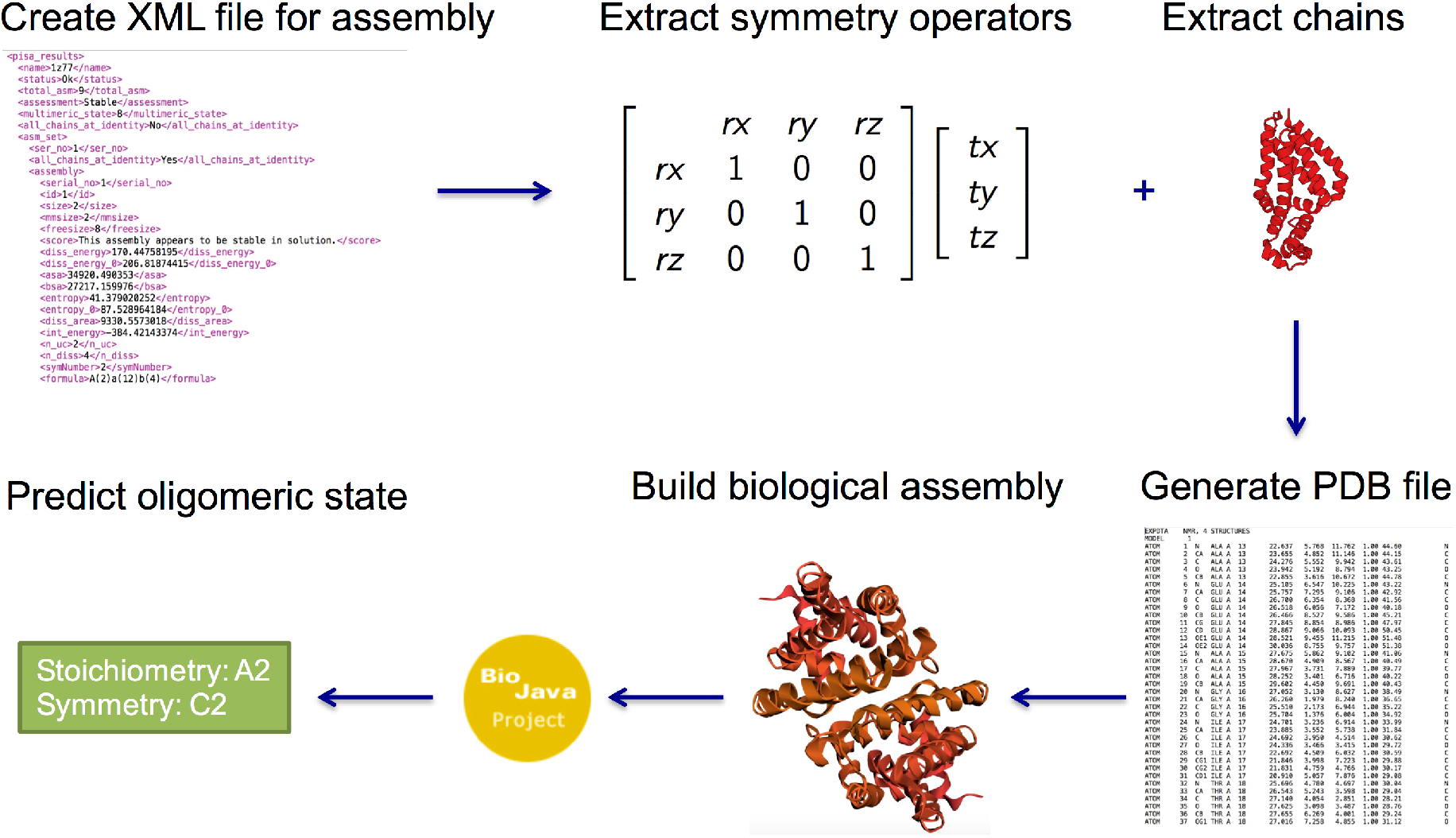
Overview of the PISA annotation procedure. An XML file is created using PISA, the file is parsed to extract symmetry operators and corresponding chains and BioJava is used to assign stoichiometry and symmetry.

### EPPIC Prediction

To distinguish intermolecular contacts stabilizing biologically relevant oligomeric states from simple crystal contacts, an evolutionary-based classifier (EPPIC; http://www.eppic-web.org) was developed by Duarte et al. [5]. This method uses a geometric measure, number of interfacial residues, and evolutionary features to classify interfaces as biological versus crystal. Unlike other evolutionary-based methods, EPPIC uses only close homologs with >60% sequence identity to ensure homologs share high degrees of quaternary structure similarity. This approach achieved ~89% accuracy with the Ponstingl et al. [23] benchmark data. The new version of EPPIC (3.0.1) uses the pairwise interface classifications from version 2, combining them to come up with quaternary structure predictions [39]. The crystal lattice is represented as a periodic graph from which the different valid assemblies (those with point group symmetry) can be enumerated. Based on the pairwise scores the software can then decide which of the viable assemblies is the most likely quaternary structure in solution. We obtained early access to the results from the authors. Output results for the whole PDB archive were provided in XML format, containing quaternary structure predictions for each entry in the PDB. We parsed these XML files to extract the respective stoichiometry and symmetry information for PDB entries.

### Consensus Result Approach

Predictions from sequence clustering, text mining, PISA, and EPPIC, were combined and a majority vote rule was applied to arrive at a consensus result. In the following cases, predictions from individual methods were excluded:

- Sequence clustering: insufficient homologous proteins comprising the cluster (at least 3 structures for 70% sequence identity).
- Text mining: no full-text publication available or no clear information regarding quaternary structure therein.
- PISA: indeterminate predictions in “gray” region.

### Web-Tool Development

To make this approach accessible, we developed a user-friendly, easy-to-use web-based tool using R, JavaScript, jQuery, CSS, and HTML (Fig 8). The tool was predominantly constructed using R software [40]. Publications were downloaded either in PDF or XML format, and XML [41], Rcurl [42], tm [43], NLP [44], openNLP [45], and stringr [46] packages were used to convert PDF files to plain text files, to parse XML files, to extract sentences, and to split words. The FeatureHashing package [47] was used to map features to the hash table [29]. Machine learning algorithms were trained and tested using the caret package [48]. Data tables were built using the DT package [49], and the shiny package [50] was used to create an interactive web-based application.

**Fig 8.**
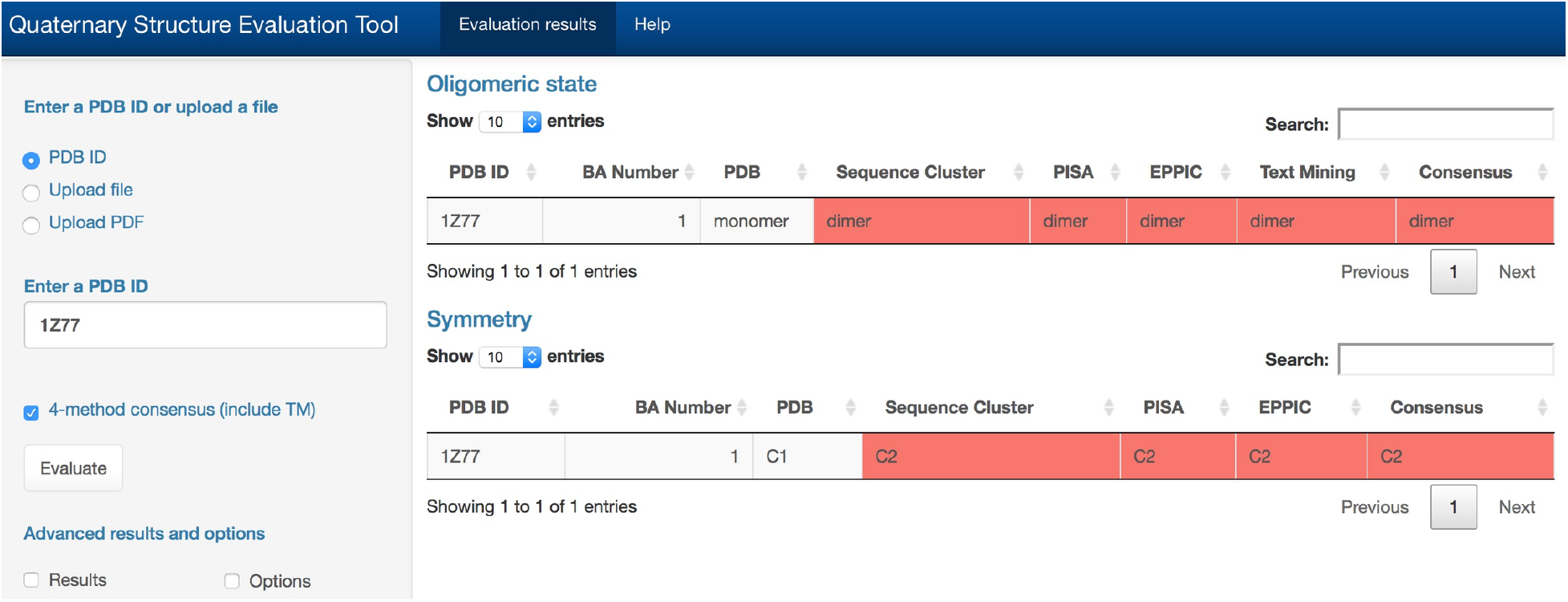
Evaluation of the quaternary structure prediction of PDB entry 1Z77. All methods (SC: sequence clustering, TM: text mining, PISA, EPPIC) are selected for the evaluation process. The output includes two parts: oligomeric state and symmetry. In the oligomeric state table, it states that 1Z77 (first column: PDB ID) has one oligomeric state prediction (second column: BA Number), and it is annotated as a monomeric structure in the PDB (third column: PDB). However, according to SC (fourth column: Sequence clustering), PISA (fifth column), EPPIC (sixth column), and TM (seventh column: Text Mining), 1Z77 is a dimeric protein. Therefore, our consensus result (eighth column) states that 1Z77 is a dimer. In the symmetry table one quaternary structure prediction is shown (second column: BA Number), having C1 symmetry (third column: PDB). However, according to the SC (fourth columns: Sequence cluster), PISA (fifth column) and EPPIC (sixth column), 1Z77 has a C2 symmetry. Therefore our consensus result (seventh column) states that 1Z77 has a C2 symmetry. Red denotes divergence between current PDB annotation of oligomeric state and results provided by each of the four evaluation methods.

The new web tool offers a wide range of methods for evaluating oligomeric states in the PDB, including

i. Determination of a representative oligomeric state for a given certain sequence identity threshold using the sequence clustering approach with consistency scoring.
ii. Generate a PISA oligomeric state prediction for a given structure, rebuild the quaternary structure, and assign stoichiometry and symmetry using BioJava.
iii. Generate a EPPIC oligomeric state prediction for a given structure, including stoichiometry and symmetry.
iv. Extract oligomeric state information with experimental evidence from any publication describing a crystal structure of a protein.

The tool has a simple user interface, requiring only four character PDB IDs as input and a single mouse click to launch the calculation. Users can either enter a single PDB entry or upload a *.txt* file, which includes multiple PDB IDs, using the option in the tool for processing multiple PDB entries. Users can also upload a PDF version of a paper within the text mining module.

## Results

### Benchmark Dataset Results

We applied sequence clustering, text mining, PISA, and EPPIC to our 543 structure benchmark dataset to test the performance of each approach. Then, we aggregated all available results to arrive at a consensus result. S3 Table lists the individual predictions and S4 Table summarizes the results for individual methods and the consensus predictions. Accuracy rates for individual methods ranged between 46% and 81%. We achieved 86% accuracy rate using the consensus approach (Fig 9).

**Fig 9.**
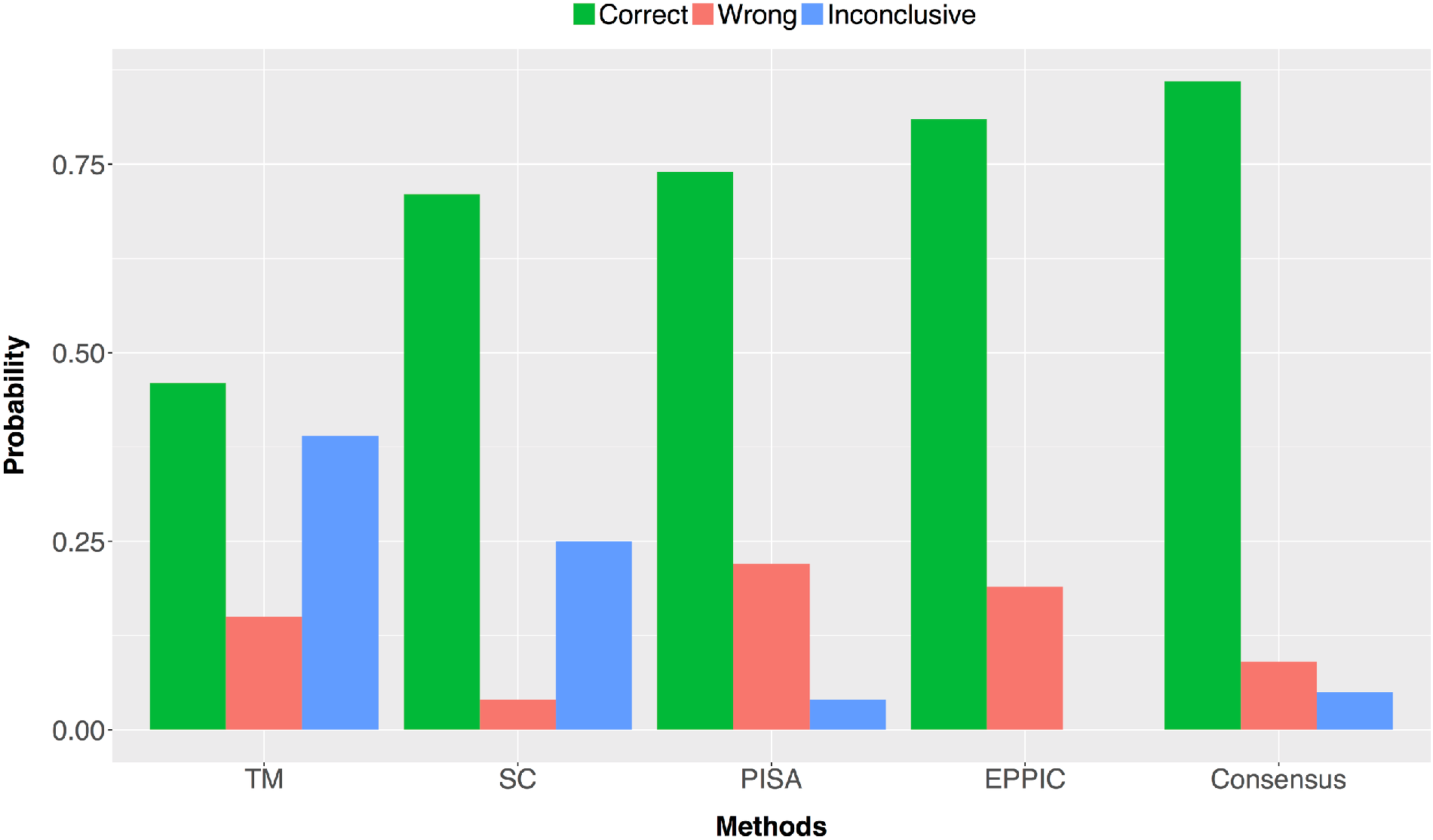
Overall performance results for the benchmark dataset using four different methods and consensus approach. The number of correct predictions for method-agreements are summarized in Fig 10A. Four-method agreement correctly predicted the oligomeric states of 122 structures. The combination of SC, PISA and EPPIC (i.e. three-method agreement) correctly predicted 149 cases. In two-method agreements, PISA and EPPIC correctly predicted 40 cases, while SC and EPPIC correctly predicted 31 cases, and SC and PISA predicted 21 cases correctly. When we checked number of incorrect predictions (Fig 10B), all methods incorrectly predicted only 1 structure. The most incorrect results occurred between PISA and EPPIC methods with 34 cases. Three methods, TM, PISA, and EPPIC, and two methods, TM and EPPIC, incorrectly predicted. Finally, inconclusive results (Fig 10C) mostly occurred between TM and SC methods with 40 cases.

**Fig 10.**
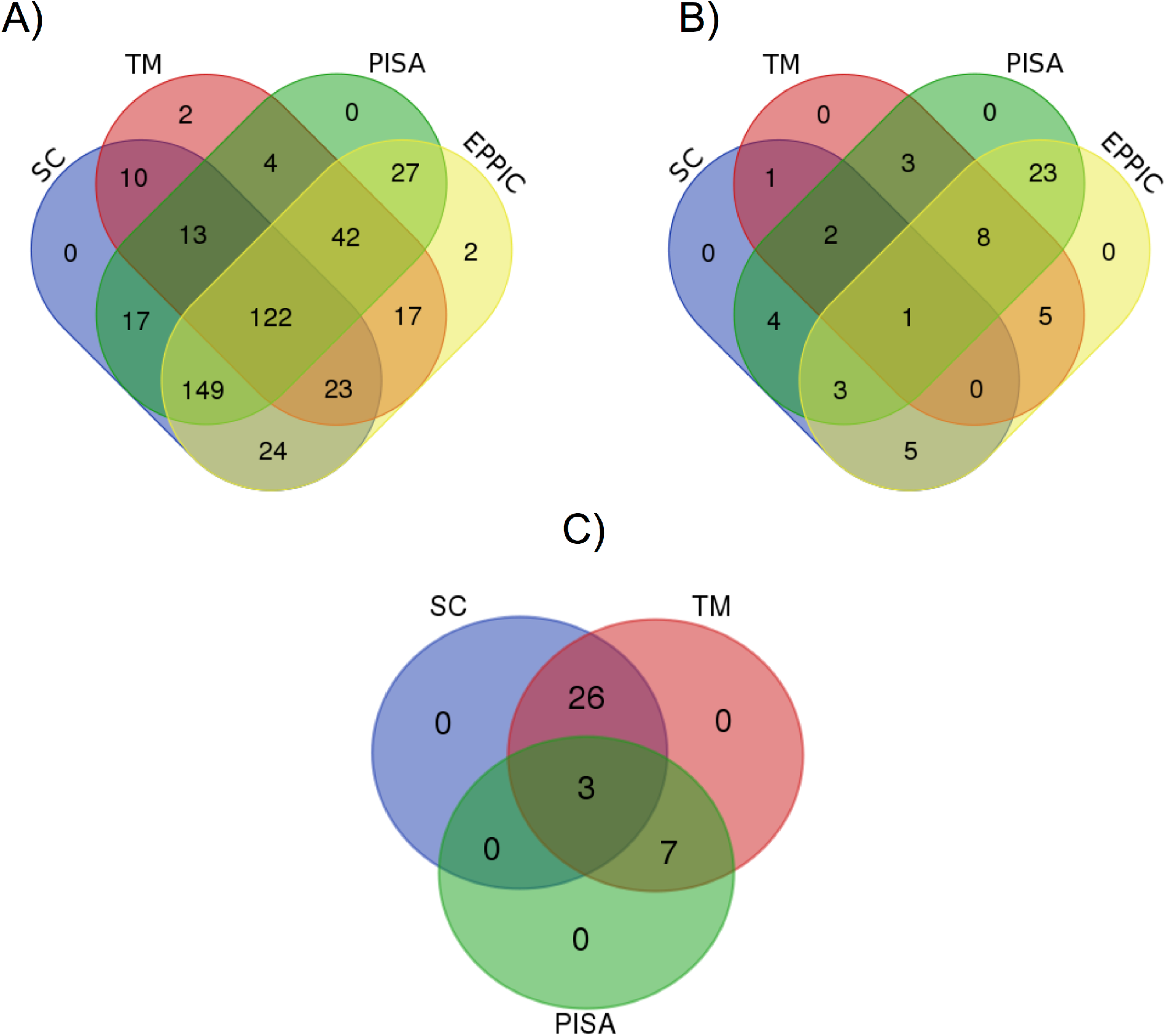
Prediction agreement of the consensus approach. A) Correct prediction agreements between methods. B) Incorrect prediction agreements between methods. C) Inconclusive prediction agreements between methods.

Since TM gives high inconlusive results, we also provide a 3-method consensus based on SC, PISA and EPPIC results. After exclusion of TM, the accuracy rate decreased to 81%, while incorrect rate increased to 11% and inconclusive rate increased to 8%. Thus, even though its highly inconclusive nature, TM provides useful insights regarding correct quaternary structure information.

## Discussion

The oligomeric state (or states) of a protein represents the essential biological unit(s) that carries out a biological function in a living organism [4]. It is, therefore, crucial to determine with high reliability the biologically-relevant oligomerization state(s) of a macromolecule. This problem has been extensively studied and the scientific literature abounds with various methods and approaches. An extensive analysis of the problem can be found in Capitani et al. [20], wherein the authors reviewed main concepts of different approaches, including thermodynamic estimation of interface stability, evolutionary approaches and interface co-occurrence across different crystal forms.

The difficulty of the protein interface classification problem has led to incomplete or ambiguous experimental data, which in turn has resulted in incorrect annotations of the oligomerization states of macromolecules in the PDB. Another source of error are the simple mistakes during the deposition process. For example, authors sometimes simply identify the quaternary structure of the asydmmetric unit. To the best of our knowledge, there are only two studies in the literature, which have investigated quaternary structure annotation errors in the PDB [21, 22].

In this study, we focused on two main issues: (i) detection of the incorrectly annotated biological macromolecules in the PDB and (ii) assignment of the most probable quaternary structures for those incorrect structures. We used two of the most popular and widely used methods to evaluate the quaternary structures of macromolecules, PISA and EPPIC. To investigate author annotations, we provided a text mining approach for searching and extracting correct quaternary structure information from primary papers of the macromolecules. In addition, we utilized homology, by looking at consensus within sequence clusters, for detection of incorrect quaternary structure annotations in the PDB. Finally, we obtained quaternary structure predictions from each method, including oligomeric state and symmetry information, and came up with a consensus result for each entry in the PDB. The consensus result approach, which uses the method agreements, has improved the prediction reliability. The number of correct predictions is increased, while both the number of incorrect predictions and the number of inconclusive predictions are decreased. The best combination of the methods is found as the three methods combination of Sequence Cluster, PISA and EPPIC, which correctly predicted 149 cases together. On the other hand, the four method agreement correctly predicted 122 quaternary structures. On the other hand, the four method agreement correctly predicted 110 quaternary structures. The reason of the lower number of correct predictions in the four method agreements is the high rate of inconclusive results in the text mining approach, which is almost 46%.

The most important issue in the text mining method is the difficulty in the full-text publication access. Even though, 42% of the primary articles in the PDB are open-access publications, it is hard to get these publications using the PMC database or publishers’ websites for text mining, because these services have their own restrictions based on their respective policies. PMC does provide a service for bulk downloading of the publications for an open-access subset. However, this subset only covers 1/26 of all articles in PubMed, and that makes it impractical for text mining purposes. Some publishers, such as Elsevier and Wiley, provide their own text mining services through a specific API (application program interface). However, these services are not stable and not working all the time and more importantly they are not allowing bulk access to many papers. In addition, they do not provide all the publications through these services; instead they only provide a small subset of them. We found CrossRef TDM services as the most useful service for text mining purposes. They provide full-text links from various publishers in a user friendly and easy-to-access way. However, since some publishers only share a small part of their paper repositories, we are only able to reach 30% of the primary articles of the PDB entries.

In addition to the 4-method consensus results, we also provided a 3-method consensus based on SC, PISA and EPPIC results by excluding TM. The 3-method consensus has the advantage that it can be applied to most structures in the PDB. On the other hand, TM results are useful to guide the user/biocurator to the specific papers/sentences that contain statements about the experimental evidence, as well as a resource to verify the author annotations. Therefore, we provided 3-method consensus as well as 4-method consensus in our web-tool.

## Conclusion

Determination of a 3D macromolecular structure is crucial to understand the fundamental mechanisms of biological processes, such as enzymatic reactions, ligand binding, or signalling. It is also important to reveal the underlying mechanisms of diseases, such as genetic variations. Furthermore, the 3D structures of macromolecules are vital for drug design and development studies, especially in structure-based drug design. Because of these reasons, the correct quaternary structure has a critical importance.

In this study, we developed a consensus approach by aggregating predictions from three and four different methods in order to detect incorrect quaternary structures in the PDB and assign the most likely quaternary structures for the possibly incorrect annotations. For this task, we first benefited from homology to cluster similar PDB entries based on a certain sequence identity threshold and to predict a representative quaternary structure for the cluster through a consistency score calculation. Second, we searched through the primary articles associated with the PDB entries to validate the author deposition by extracting oligomeric state and experimental evidence information using a text mining approach. Moreover, we used the PISA and EPPIC to predict the quaternary structures. Then, we combined predictions from different approaches to achieve a consensus result and to predict the most probable quaternary structures for the possibly incorrect structures. To test the performance of our consensus approach, we created a benchmark dataset using Ponstingl et al. [23], Bahadur et al. [24], Bahadur et al. [25] and Duarte et al. [5] datasets. Our consensus approach outperformed single methods and achieved 86% correct, 9% incorrect, and 5% inconclusive predictions, respectively. Therefore, our method provides more reliable evaluation than any single approach. Finally, we developed a web-based tool in order to make this approach usable for researchers in the field and PDB Biocuators.

## Availabilty

The tool is freely available through http://quatstruct.rcsb.org. All source code is available on Github repository at https://github.com/selcukorkmaz/BET. This tool will be updated regularly to include the future quaternary structures in the PDB.

## Acknowledgments

We thank Spencer Bliven, Guido Capitani, Anthony Bradley, and Alex Rose for their suggestions and useful discussions, and Cole H. Christie and Chris Randle for making the web-tool accessible. We also thank Eugene Krissinel for explaining usage of the PISA software.

---

**S1 Table. Keywords list for searching quaternary structure related sentences.**

**S2 Table. Keywords list for searching experimental evidence related sentences.**

**S3 Table. Predictions on benchmark sets using 4 methods and consensus results.**

**S4 Table. Summary of prediction results for benchmark sets.**

## References

1 Berman HM, Westbrook J, Feng Z, Gilliland G, Bhat TN, Weissig H, et al. The Protein Data Bank. Nucleic Acids Res. 2000;28(1):235–242. doi: 10.1093/nar/28.1.235.

2 Berman H, Henrick K, Nakamura H. Announcing the worldwide Protein Data Bank. Nat Struct Biol. 2003;10(12):980–980. doi: 10.1038/nsb1203-980.

3 Goodsell DS, Olson AJ. Structural symmetry and protein function. Annu Rev Biophys Biomol Struct.. 2000;29(1):105–153. doi: 10.1146/annurev.biophys.29.1.105. PMID: 10940245.

4 Krissinel E, Henrick K. Inference of macromolecular assemblies from crystalline state. J Mol Biol. 2007;372(3):774–797. doi: 10.1016/j.jmb.2007.05.022. PMID: 17681537.

5 Duarte JM, Srebniak A, Scharer MA, Capitani G. Protein interface classification by evolutionary analysis. BMC Bioinformatics. 2012;13(334):1–16. doi: Artn 334 10.1186/1471-2105-13-334.

6 Henrick K, Thornton JM. PQS: a protein quaternary structure file server. Trends Biochem Sci. 1998;23(9):358–361. PMID: 9787643.

7 Rose PW, Prlic A, Altunkaya A, Bi C, Bradley AR, Christie CH, et al. The RCSB protein data bank: integrative view of protein, gene and 3D structural information. Nucleic Acids Res. 2016;45(D1);D271–D281. doi: 10.1093/nar/gkw1000. PMID: 27794042.

8 Winzor DJ. Analytical exclusion chromatography. J Biochem Biophys Methods. 2003;56(1):15–52. PMID: 12834967.

9 Lebowitz J, Lewis MS, Schuck P. Modern analytical ultracentrifugation in protein science: a tutorial review. Protein Sci. 2002;11(9):2067–2079. doi: 10.1110/ps.0207702. PMID: 12192063.

10 Fasshauer D, Otto H, Eliason WK, Jahn R, Brunger AT. Structural changes are associated with soluble N-ethylmaleimide-sensitive fusion protein attachment protein receptor complex formation. J Biol Chem. 1997;272(44):28036–28041. doi: 10.1074/jbc.272.44.28036.

11 Takamori S, Holt M, Stenius K, Lemke EA, Gronborg M, Riedel D, et al. Molecular anatomy of a trafficking organelle. Cell. 2006;127(4):831–846. doi: 10.1016/j.cell.2006.10.030. PMID: 17110340.

12 Myers-Turnbull D, Bliven SE, Rose PW, Aziz ZK, Youkharibache P, Bourne PE, et al. Systematic Detection of Internal Symmetry in Proteins Using CE-Symm. J Mol Biol. 2014;426(11):2255–2268. doi: 10.1016/j.jmb.2014.03.010.

13 Lee J, Blaber M. Experimental support for the evolution of symmetric protein architecture from a simple peptide motif. P Natl Acad Sci USA. 2011;108(1):126–130. doi: 10.1073/pnas.1015032108.

14 Wolynes PG, Luthey-Schulten Z, Onuchic JN. Fast-folding eriments and the topography of protein folding energy landscapes. Chem Biol. 1996;3(6):425–432.

15 Waldrop GL. The role of symmetry in the regulation of bacterial carboxyltransferase. Biomol Concepts. 2011;2(1–2):47–52.

16 Changeux JP, Edelstein SJ. Allosteric mechanisms of signal transduction. Science. 2005;308(5727):1424–1428. doi: 10.1126/science.1108595. PMID: 15933191.

17 Monod J, Wyman J, Changeux JP. On the Nature of Allosteric Transitions: A Plausible Model. J Mol Biol. 1965;12:88–118. PMID: 14343300.

18 Marsh JA, Rees HA, Ahnert SE, Teichmann SA. Structural and evolutionary versatility in protein complexes with uneven stoichiometry. Nat Commun. 2015;6:6394. doi: 10.1038/ncomms7394. PMID: 25775164.

19 Levy ED, Boeri Erba E, Robinson CV, Teichmann SA. Assembly reflects evolution of protein complexes. Nature. 2008;453(7199):1262–1265. doi: 10.1038/nature06942. PMID: 18563089.

20 Capitani G, Duarte JM, Baskaran K, Bliven S, Somody JC. Understanding the fabric of protein crystals: computational classification of biological interfaces and crystal contacts. Bioinformatics. 2016;32(4):481–489. doi: 10.1093/bioinformatics/btv622. PMID: 26508758.

21 Levy ED. PiQSi: protein quaternary structure investigation. Structure. 2007;15(11):1364–1367. doi: 10.1016/j.str.2007.09.019. PMID: 17997962.

22 Baskaran K, Duarte JM, Biyani N, Bliven S, Capitani G. A PDB-wide, evolution-based assessment of protein-protein interfaces. BMC Struct Biol. 2014;14(1):1–11. doi: 10.1186/s12900-014-0022-0.

23 Ponstingl H, Kabir T, Thornton JM. Automatic inference of protein quaternary structure from crystals. J Appl Crystallogr. 2003;36(5):1116–1122. doi: 10.1107/S0021889803012421.

24 Bahadur RP, Chakrabarti P, Rodier F, Janin J. Dissecting subunit interfaces in homodimeric proteins. Proteins. 2003;53(3):708–719. doi: 10.1002/prot.10461.

25 Bahadur RP, Chakrabarti P, Rodier F, Janin J. A dissection of specific and non-specific protein - Protein interfaces. J Mol Biol. 2004;336(4):943–955. doi: 10.1016/j.jmb.2003.12.073.

26 Dondoshansky I, Wolf Y. Blastclust (ncbi software development toolkit). NCBI, Bethesda, Md 2002.

27 Altschul SF, Gish W, Miller W, Myers EW, Lipman DJ. Basic local alignment search tool. J Mol Biol. 1990;215(3):403–410. doi: 10.1016/S0022-2836(05)80360-2. PMID: 2231712.

28 Leskovec J, Rajaraman A, Ullman JD. Mining of massive datasets. United Kingdom: Cambridge University Press; 2014.

29 Weinberger K, Dasgupta A, Langford J, Smola A, Attenberg J. Feature hashing for large scale multitask learning. Proceedings of the 26th Annual International Conference on Machine Learning. 2009:1113–1120.

30 Vapnik V. The nature of statistical learning theory. New York: Springer 2000.

31 Friedman J, Hastie T, Tibshirani R. Additive logistic regression: A statistical view of boosting - Rejoinder. Ann Stat. 2000;28(2):400–407.

32 Freund Y. Boosting a Weak Learning Algorithm by Majority. Inf Comput. 1995;121(2):256–285. doi: 10.1006/inco.1995.1136.

33 Freund Y, Schapire RE. A decision-theoretic generalization of on-line learning and an application to boosting. J Comput Syst Sci. 1997;55(1):119–139. doi: 10.1006/jcss.1997.1504.

34 Schapire RE, Singer Y. Improved boosting algorithms using confidence-rated predictions. Mach Learn. 1999;37(3):297–336. doi: 10.1023/A:1007614523901.

35 Krissinel E. Stock-based detection of protein oligomeric states in jsPISA. Nucleic Acids Res. 2015;43(W1):W314–W319. doi: 10.1093/nar/gkv314.

36 Gopavajhula VR, Chaitanya KV, Khan PAA, Shaik JP, Reddy PN, Alanazi M. Modeling and analysis of soybean (Glycine max. L) Cu/Zn, Mn and Fe superoxide dismutases. Genet Mol Biol. 2013;36(2):225–236.

37 Krissinel E. Macromolecular complexes in crystals and solutions. Acta Crystallogr D. 2011;67(4):376–385. doi: 10.1107/S0907444911007232.

38 Prlic A, Yates A, Bliven SE, Rose PW, Jacobsen J, Troshin PV, et al. BioJava: an open-source framework for bioinformatics in 2012. Bioinformatics. 2012;28(20):2693–2695. doi: 10.1093/bioinformatics/bts494.

39 Duarte JM, Bliven S, Liafita A, Parker A, Capitani G, editors. Automated finding of assemblies in protein crystal structures. 3D SIG - Structural Bioinformatics and Computational Biophysics 2016; Orlando, Florida: F1000Research 2016.

40 R Core Team. R: A language and environment for statistical computing. R Foundation for Statistical Computing. Vienna, Austria URL https://wwwr-projectorg/. 2016.

41 Lang DT. XML: Tools for Parsing and Generating XML Within R and S-Plus. R Package Version 398–142016. p. https://cran.r-project.org/web/packages/XML/index.html.

42 Lang DT. RCurl: General Network (HTTP/FTP/…) Client Interface for R. R Package Version 195–482016. p. https://cran.r-proiect.org/web/packages/RCurl/index.html.

43 Feinerer I, Hornik K. tm: Text Mining Package. R Package Version 06–22015. p. https://cran.r-project.org/web/packages/tm/index.html.

44 Hornik K. NLP: Natural Language Processing Infrastructure. R Package Version 01–92016. p. https://cran.r-project.org/web/packages/NLP/index.html.

45 Hornik K. openNLP: Apache OpenNLP Tools Interface. R Package Version 02–62016. p. https://cran.r-project.org/web/packages/openNLP/index.html.

46 Wickam H. stringr: Simple, Consistent Wrappers for Common String Operations. R Package Version 1002015. p. https://cran.r-project.org/web/packages/stringr/index.html.

47 Wu W. FeatureHashing: Creates a Model Matrix via Feature Hashing with a Formula Interface. R Package Version 09112015. p. https://cran.r-project.org/web/packages/FeatureHashing/index.html.

48 Kuhn M. Building Predictive Models in R Using the caret Package. J Stat Softw2008. p. 1–26.

49 Xie Y, Cheng J, Allaire JJ, Reavis B, Gersen L, Szopka B. DT: A Wrapper of the JavaScript Library 'DataTables'. Available at https://cranr-projectorg/web/packages/DT/. 2016.

50 Chang W, Cheng J, Allaire JJ, Xie Y, McPherson J. shiny: Web Application Framework for R. R Package Version 01312016. p. https://cran.r-project.org/web/packages/shiny/index.html.

